# Targeted delivery of interferon-gamma by synNotch T cells sensitizes neuroblastoma cells to T cell-mediated killing

**DOI:** 10.1101/2022.06.21.496901

**Authors:** Fiorella Iglesias, Siani Weston, Stephanie V. Avila, Holly Zhou, Sara Yousef, Mark Fluchel, Erin Morales, Michael L. Olsen, Sabarinath V. Radhakrishnan, Tim Luetkens

## Abstract

Downregulation of HLA is one of the most common tumor escape mechanisms by enabling tumors to persist in the presence of tumor-reactive T cells. HLA loss is particularly common in children with high-risk neuroblastoma, who have a 50% long-term survival despite dose-intensive regimens. We have now developed an approach for the targeted induction of HLA to restore sensitivity of neuroblastoma cells to T cell-mediated killing. Using synNotch technology, we have generated T cells that, upon binding of the neuroblastoma surface antigen GD2, secrete IFN-γ without conferring direct cytotoxic activity (snGD2i). Treatment with snGD2i cells induces high and durable expression of HLA on neuroblastoma cells *in vitro* and *in vivo* and restores sensitivity to TCR-transgenic T cells targeting neuroblastoma-specific antigens. In contrast, treatment does not lead to upregulation of immune checkpoints or systemically increased levels of IFN-γ. Targeted delivery of IFN-γ using snGD2i cells is a promising new strategy to address immune escape in neuroblastoma.

**STATEMENT OF SIGNIFICANCE:** HLA loss remains one of the most common and unsolved immune escape mechanisms in cancer cells. We have now developed a cell-based approach for the targeted upregulation of HLA on tumor cells, which efficiently sensitizes the malignant cells to killing by tumor-reactive T cells.

## INTRODUCTION

Cancer-specific T cells are able to control tumor growth through the recognition of tumor-associated antigens presented by human leukocyte antigen (HLA) proteins. However, an established mechanism of immune evasion in pediatric malignancies, such as neuroblastoma, is the downregulation of HLA class I molecules ^1^. This not only prevents the patient’s own cancer-specific T cells from recognizing the malignant cells but also reduces the effectiveness of T cell receptor (TCR)-based immunotherapies and immune checkpoint inhibitors ^2^. Despite the high prevalence of this phenomenon, to date no clinically-proven approaches exist to efficiently restore HLA expression on tumor cells.

Neuroblastoma is a sympathetic nervous system malignancy that arises from neural crest cells and is responsible for more than 10% of deaths associated with cancer in children under 15 years of age ^3^. Five-year event-free survival for high-risk neuroblastoma patients is only 50%, even following intensive multi-modal treatment demonstrating the urgent need for novel therapeutic approaches ^4^. We hypothesize that restoring HLA expression on neuroblastoma cells will facilitate their killing by pre-existing tumor-reactive patient T cells analogous to restoring anti-tumor immunity using checkpoint inhibitors.

In addition to re-activating such pre-existing anti-tumor immunity, the adoptive transfer of receptor-transgenic T cells has proven highly successful in the treatment of various hematologic pediatric malignancies, such as ALL ^5^. CAR T cell approaches targeting the surface antigens GD2 ^6^, B7H3 ^7^, L1CAM ^8^, GPC2 ^9^, NCAM ^10^, and ALK ^11^ are currently being explored as novel therapeutics in neuroblastoma ^12^. However, unintended targeting of healthy cells expressing the respective target antigen still raises concerns when using these approaches ^13^ and modest responses and short persistence remain ongoing issues ^14^.

Solid cancers, including neuroblastoma, have been shown to frequently express various highly specific intracellular tumor antigens, that, while generally inaccessible to conventional CAR T cell approaches, can be targeted using TCR-transgenic T cells ^15^. While TCR T cells showed preclinical promise for the treatment of neuroblastoma ^16,17^, these therapies are unlikely to be effective clinically due to the pronounced HLA loss. Here, we develop a novel cell-based strategy for the targeted restoration of HLA expression on neuroblastoma cells to facilitate T cell-mediated anti-tumor immunity as a novel therapeutic approach for the treatment of patients with neuroblastoma.

## RESULTS

### HLA class I expression is downregulated in poorly differentiated/MYCN amplified neuroblastoma

One of the most common immune escape mechanisms of cancer cells is the downregulation of HLA class I and neuroblastoma shows some of the most frequent and severe HLA loss ^18^. In several cancers, HLA loss has been shown to be associated with advanced stage and less differentiated tumors ^2^. We therefore first determined HLA class I expression on primary tumor samples from patients with different types of neuroblastoma using immunohistochemistry (Suppl. Fig. 1). We observed strong HLA class I expression in well-differentiated neuroblastoma tumors (Fig. 1A). However, all 6/6 analyzed patients with poorly differentiated tumors, for whom novel therapeutic strategies are most urgently needed, showed loss of HLA class I expression on their tumor cells. Analyzing neuroblastoma cell lines by flow cytometry, we further observed substantially lower levels of surface HLA class I expression on more aggressive MYCN-amplified tumor cell lines (Fig. 1B). Restoration of HLA class I may represent a particularly promising approach to facilitate T cell-mediated anti-tumor immunity especially in patients with poorly differentiated neuroblastoma.

**Figure 1:**
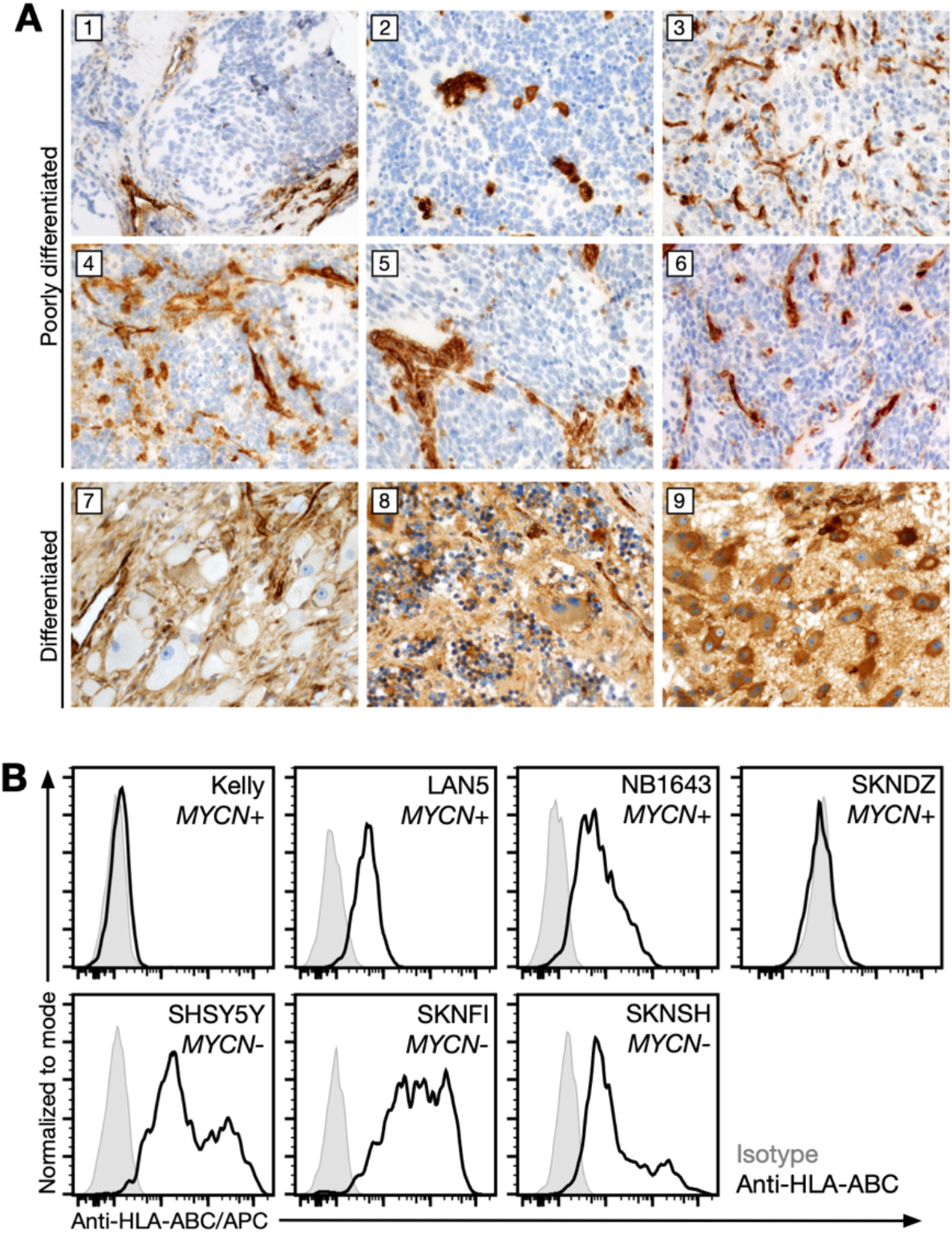
HLA class I is downregulated in poorly differentiated neuroblastoma. **(A)** HLA-ABC immunohistochemistry staining of formalin-fixed paraffin-embedded tissue from previously untreated neuroblastoma tumors. 1-6: In poorly differentiated neuroblastomas (low-intermediate MKI), the neuroblastic cells show negative HLA staining (original magnification 100x). 7-8: In ganglioneuroblastoma, intermixed subtype, the tumor cells show variable but increased HLA expression with cytoplasmic and membrane staining in tumor cells with ganglionic features (original magnification 200x). 9: In differentiating neuroblastoma, the differentiating neuroblastic cells show increased HLA expression with cytoplasmic and membrane staining in tumor cells with ganglionic features (original magnification 200x). **(B)** Expression of HLA-ABC in MYCN-amplified (MYCN+) and MYCN-wildtype (MYCN-) neuroblastoma cell lines as determined by flow cytometry after staining with an HLA-ABC-specific antibody or an isotype control antibody.

### Recombinant IFN-γ and T cells constitutively expressing IFN-γ induce HLA class I expression on neuroblastoma cells

It has previously been shown that IFN-γ can increase the expression of HLA on various cell types including some neuroblastoma tumor cells ^19,20^. Treating neuroblastoma cells with recombinant IFN-γ, we indeed observed strong HLA class I upregulation on the cell surface of all 3/3 analyzed MYCN-amplified neuroblastoma cell lines (Fig. 2A), and a substantial increase in HLA-A2 mRNA expression in the HLA-A2^+^ (Suppl. Fig. 2) neuroblastoma cell line SKNDZ (Fig. 2B).

**Figure 2:**
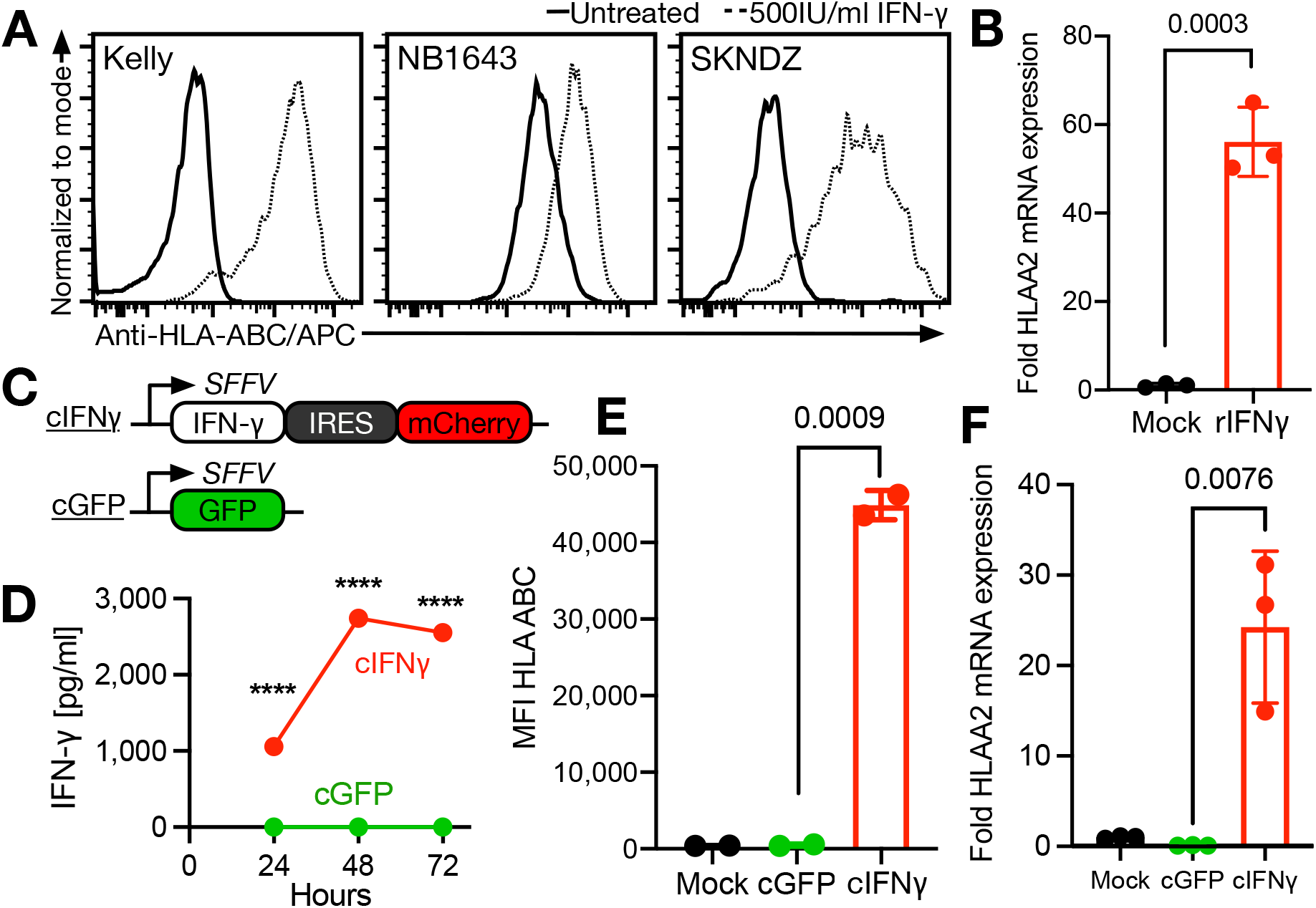
HLA class I is induced in neuroblastoma cells by recombinant IFN-γ and by T cells engineered to constitutively express IFN-γ. **(A)** Expression of HLA class I (HLA-ABC) by MYCN+ neuroblastoma cell lines with or without prior recombinant IFN-γ treatment for 48h as determined by flow cytometry. Plots show a representative result from 4 independent experiments. **(B)** Change in HLA-A2 mRNA expression in neuroblastoma cell line SKNDZ after treatment with 500IU/ml recombinant IFN-γ (rIFNγ) for 48h as determined by quantitative RT-PCR. Data represent mean ± SD from 3 technical replicates. Statistical significance of differences between treatment conditions was determined by two-sided Student’s *t* test. **(C)** Schema of constructs used to engineer J76 T cells to constitutively express IFN-γ (cIFNγ) or GFP (cGFP). **(D)** Secretion of IFN-γ by engineered J76 T cells over 72h as determined by ELISA. Data represent mean ± SD from 3 technical replicates. **(E)** Surface expression of HLA class I shown as mean fluorescence intensity (MFI) on neuroblastoma cell line SKNDZ after 48h co-culture with cIFNγ or cGFP J76 T cells as determined by flow cytometry. Data represent mean ± SD from 2 technical replicates. Statistical significance of differences between treatment conditions was determined by two-sided Student’s *t* test. **(F)** Change in HLA-A2 mRNA expression in neuroblastoma cell line SKNDZ after 48h co-culture with cIFNγ or cGFP J76 T cells as determined by quantitative RT-PCR. Data represent mean ± SD from 3 technical replicates. Statistical significance of differences between treatment conditions was determined by two-sided Student’s *t* test.

Systemic injection of recombinant IFN-γ has largely been abandoned as a viable treatment strategy, as it has been shown to result in substantial toxicities and relatively low intratumoral concentrations of the cytokine ^21,22^. We therefore propose the targeted delivery of IFN-γ using transgenic T cells. To this end, we first engineered TCR-deficient J76 T cells to constitutively express IFN-γ (cIFNγ, Fig. 2C) to determine if such overexpression would result in efficient IFN-γ secretion. We found that constitutive overexpression indeed resulted in increasing levels of the cytokine in culture supernatants over time (Fig. 2D). Co-culturing cIFNγ T cells with neuroblastoma cells, we observed a significant increase in HLA class I surface expression (Fig. 2E) and HLA-A2 mRNA expression (Fig. 2F) in the neuroblastoma cells compared to co-culture with T cells constitutively expressing GFP (cGFP). These data demonstrate that HLA class I can be induced efficiently using IFN-γ and that transgenic T cells are able to produce sufficient quantities of IFN-γ to induce HLA class I on neuroblastoma cells.

### Engineered synNotch T cells specifically secrete IFN-γ in the presence of neuroblastoma cells

To avoid systemic side effects of IFN-γ treatment and to increase the local intratumoral concentration of IFN-γ, we next set out to develop a cell therapy approach to specifically deliver IFN-γ to neuroblastoma cells. To this end, we adapted the previously described synNotch system ^23,24^, in which binding of a target antigen, for example using a single-chain variable fragment (scFv), leads to cleavage of an internal Notch1 domain, which in turn releases a transcription factor that drives expression of a gene of interest (Fig. 3A). For target recognition, our receptors use a GD2-specific scFv or a CD19-specific negative control scFv. The scFvs are fused to the Notch 1 regulatory and transmembrane domains, and a heterologous transcription factor combining the yeast GAL4 DNA-binding element with the VP64 transcriptional activator. (Fig. 3B constructs I and II). In addition, these cells were transduced with a response element construct containing a PGK promotor that drives constitutive expression of a *blue fluorescent protein* (BFP) reporter and conditional expression of IFN-γ and the fluorescent reporter *mCherry* (rIFNγ, Fig. 3B construct III). The conditional response element is under the control of a minimal CMV promoter containing GAL4 upstream activation sequences (UAS) allowing binding of the GAL4-VP64 transcription factor as previously described ^25^.

**Figure 3:**
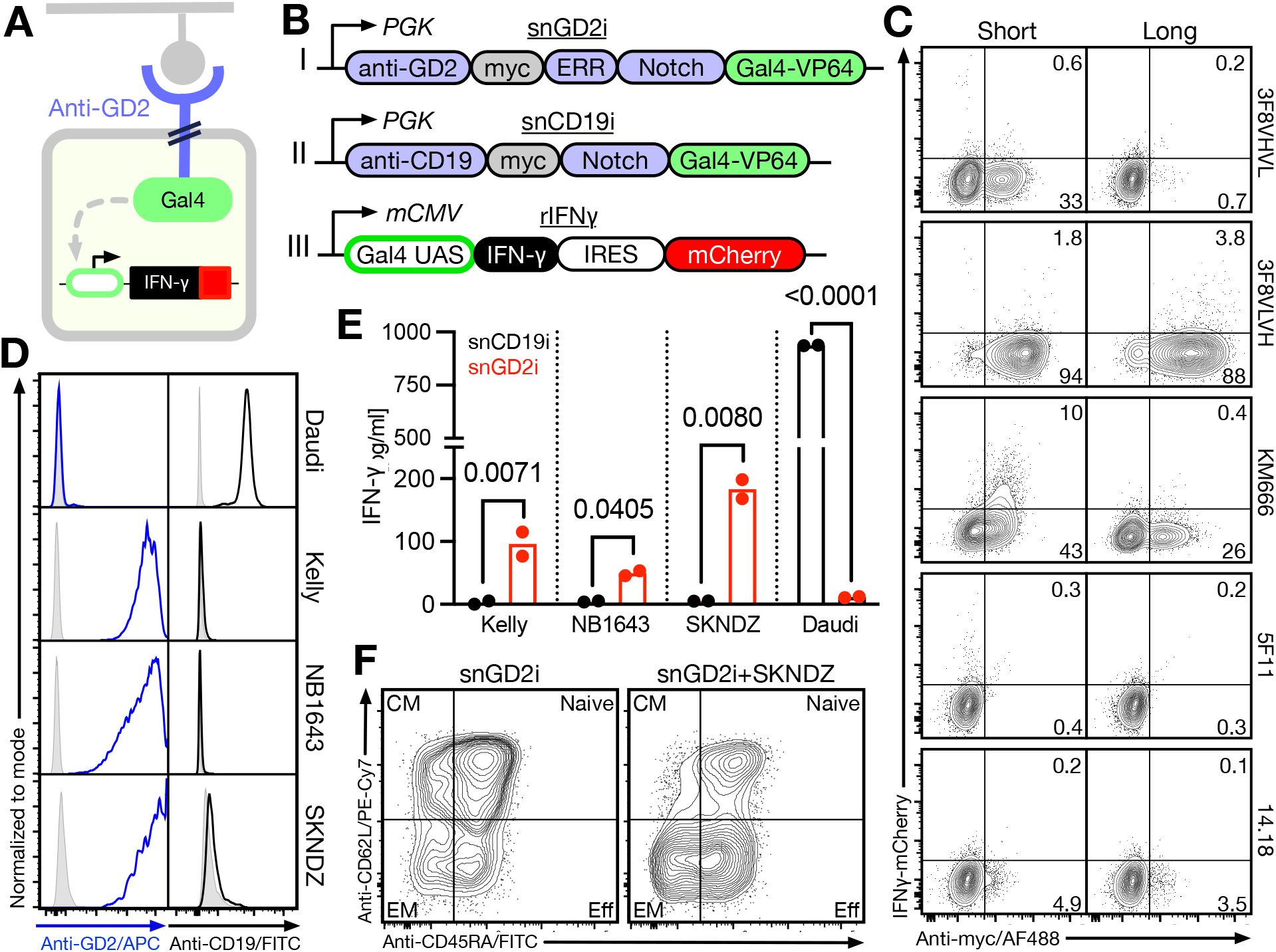
T cells engineered to express a GD2-specific synNotch receptor secrete INF−γ when co-cultured with MYCN+ neuroblastoma cells. **(A)** Schematic of a GD2-specific synNotch receptor T cell approach to induce expression of INF−γ. **(B)** Constructs used to generate synNotch T cells by combining transduction of the rINFγ construct with either the snGD2 or the snCD19 construct. **(C)** Comparison of different single chain Fv (scFv) binding domains targeting GD2 using the standard (short) core Notch1 domain or a Notch1 domain with an extended regulatory repeat region (long). Surface expression was determined using an anti-myc/AF488 antibody and baseline synNotch signaling was determined by mCherry expression from the rINFγ construct using flow cytometry. **(D)** Surface expression of GD2 and CD19 on B cell lymphoma cell line Daudi and MYCN+ neuroblastoma cell lines as determined by flow cytometry after staining with an anti-GD2/APC or an anti-CD19/FITC antibody. **(E)** Secretion of INF−γ by snGD2 and snCD19 J76 T cells during 48h co-culture with GD2^+^ neuroblastoma cell lines or a GD2^-^ lymphoma cell line as determined by ELISA. Data represent the mean of 2 technical replicates. **(F)** T cell phenotype of primary snGD2i T cells before and after co-culture with SKNDZ as determined by flow cytometry. CM= central memory; EM= effector memory; Eff= effector.

As no GD2-specific synNotch receptors have been developed previously, we first screened a set of GD2-specific scFv domains previously developed for use in conventional CAR constructs or as monoclonal antibodies. After stable transduction of J76 T cells with synNotch receptors based on those binding domains together with the response element, we determined receptor surface expression levels as well as baseline signaling, evidenced by the expression of mCherry (Figure 3C). Only 3/5 GD2 specific antibody constructs, clones 3F8 VHVL, 3F8 VLVH ^26^, and KM666 ^27^ showed surface expression on J76 cells. The synNotch receptor using the KM666-derived binding domain, however, showed substantial expression of *mCherry* in the absence of *GD2*-positive target cells indicating high basal signaling. Out of the three candidate receptors, the 3F8 construct in VL-VH orientation showed the highest surface expression in the absence of basal signaling (Figure 3C). Extending the core Notch1 domain with additional EGF repeats ^23^ further increased the surface expression of this receptor (Figure 3C), and this construct was selected for downstream assays. We next determined expression of GD2 on neuroblastoma cell lines showing HLA downregulation using flow cytometry. We found that all 3/3 analyzed neuroblastoma cell lines showed expression of GD2. Importantly, negative control lymphoma cell line Daudi showed expression of CD19 but not GD2 (Fig. 3D).

We next engineered primary human T cells to express our new GD2-synNotch receptor and carry the IFN−γ response element (snGD2i). Co-culturing snGD2i cells with the three GD2^+^CD19^-^ neuroblastoma cell lines and the GD2^-^CD19^+^ lymphoma cell lines, we observed specific secretion of IFN−γ only in the presence of neuroblastoma cell lines. Interestingly, we observed substantially higher expression of IFN−γ by CD19-specific synNotch T cells when co-cultured with the lymphoma cell line despite higher levels of GD2 on the surface of the neuroblastoma cell lines (Figure 3E). A potential explanation for this finding may be the presence of soluble GD2 produced by neuroblastoma cell lines ^28^ and we observed that substantial levels of GD2 had transferred to the surface of snGD2i cells potentially limiting their ability to produce IFN−γ over an extended period of time (Suppl. Fig. 3). Nevertheless, our data show that substantial IFN−γ levels can be achieved by snGD2i cells in the presence of neuroblastoma cells.

Finally, IFN−γ is a pleiotropic cytokine exerting various functions not only on target cells but also in T cells ^21^. We therefore explored how signaling through the synNotch receptor of snGD2i cells after binding to neuroblastoma cells shapes the T cells’ phenotype. Interestingly, we found that similar to physiological T cell activation, snGD2i cells differentiated into an effector/effector memory phenotype when co-cultured with neuroblastoma cells (Fig. 3F).

### GD2-synNotch T cells sensitize neuroblastoma cells to T cell-mediated killing

Co-culturing neuroblastoma cell line SKNDZ with snGD2i cells for 48h, we observed strong induction of HLA class I surface expression on the malignant cells (Fig. 4A). Importantly, this effect was reversed by addition of an IFN-γ blocking antibody confirming that HLA upregulation was in fact mediated by IFN-γ. Investigating whether the levels of IFN-γ secreted by snGD2i cells would be sufficient to induce HLA expression not only in neuroblastoma cells directly in contact with snGD2i cells but also bystander neuroblastoma cells, we next harvested conditioned supernatants from 48h cocultures of SKNDZ cells with snGD2i cells and treated SKNDZ cells with these conditioned supernatants for 48h. Again, we observed strong induction of HLA class I on the surface of neuroblastoma cells (Fig. 4B) as well as HLA-A2 mRNA (Fig. 4C) indicating that overall IFN-γ levels secreted were sufficient for HLA induction in bystander tumor cells without requiring immune synapse formation. HLA expression following treatment and subsequent removal of snGD2i cells remained upregulated for 8 days but showed a continued decrease over this time (Fig. 4D). We also observed a robust increase in HLA class I expression *in vivo* in large established neuroblastoma tumors treated with intratumorally injected snGD2i cells (Fig. 4E). Importantly, intratumoral delivery of snGD2i cells did not lead to systemically increased levels of IFN-γ in the same animals (Fig. 4F). IFN-γ at low levels is unlikely to have direct cytotoxic effects on tumor cells themselves. The goal of our approach is to instead augment pre-existing anti-tumor T cell responses or adoptively transferred TCR-transgenic T cells by rendering the tumor cells visible to these cells. We therefore set out to determine whether treatment with snGD2i cells would in fact sensitize neuroblastoma cells to killing by tumor specific TCR-transgenic T cells. Two well established targets for TCR T cell therapy, which had previously also been shown to be expressed in neuroblastoma, are NYESO1 and PRAME ^16,17^. While we did not observe expression of NYESO1, 3/3 cell lines expressed PRAME (Fig. 4G). To also assess killing by NYESO1-specific T cells, we transduced HLA-A2^+^ neuroblastoma cell line SKNDZ with an NYESO1 expression construct (SKNDZ-ESO). Importantly, we observed efficient killing of SKNDZ-ESO cells with both PRAME and NYESO1-specific T cells after pretreatment with snGD2i cells but not after treatment with snCD19 cells (Fig. 4H). Next, we determined the efficacy of primary snGD2 T cells together with PRAME TCR-transgenic T cells against neuroblastoma *in vivo* (Fig. 4I). Concurrent treatment with primary snGD2 but not snCD19 cells led to a significant and sustained reduction in neuroblastoma tumor size by PRAME TCR T cells (Fig. 4J), indicating that snGD2 cells augment tumor antigen-specific TCR T cell therapy *in vivo*.

**Figure 4:**
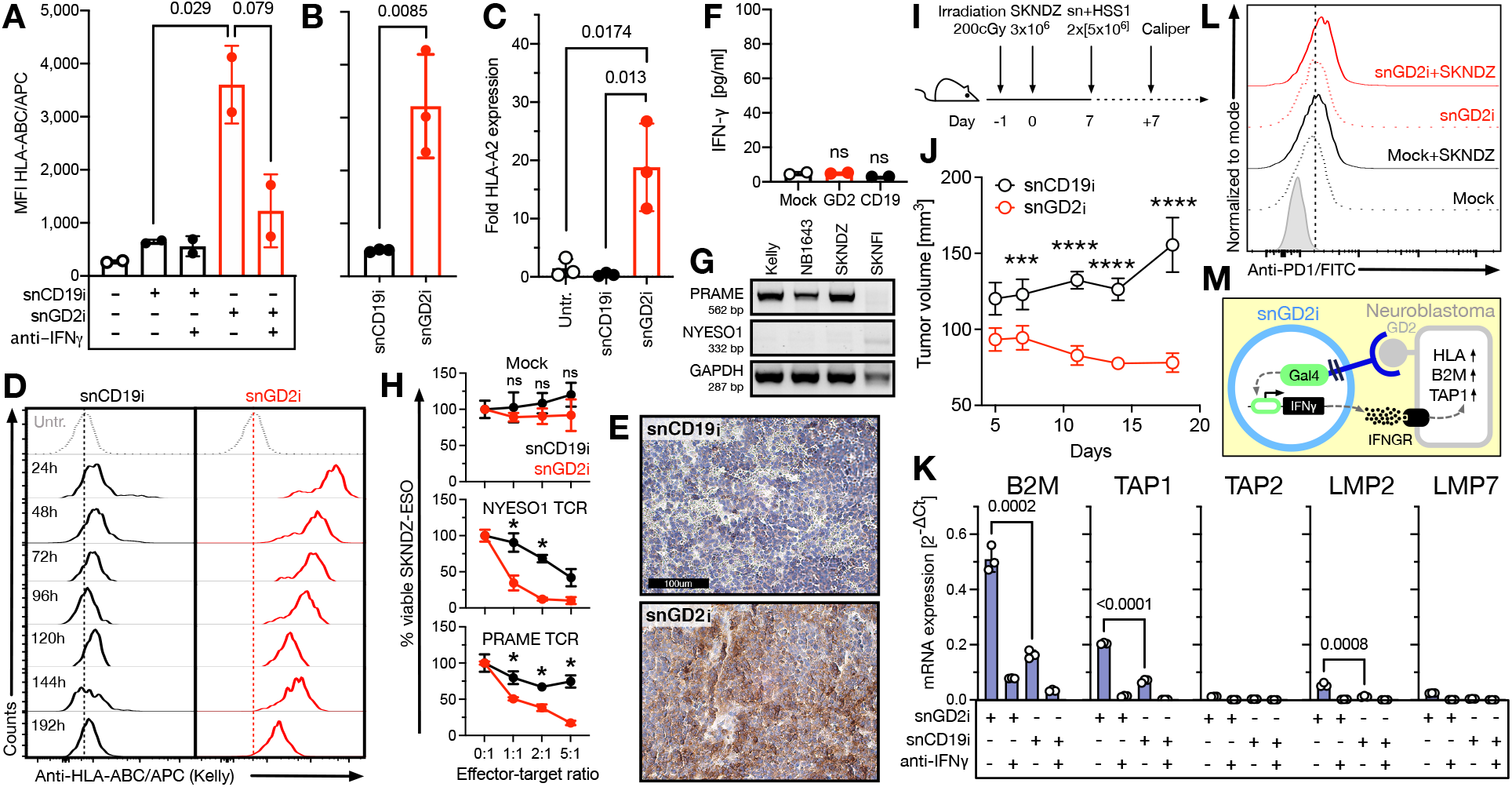
GD2-specific synNotch T cells induce HLA-ABC expression in neuroblastoma cells and augment T cell-mediated tumor killing. **(A)** Surface expression of HLA-ABC on neuroblastoma cell line SKNDZ after 48h co-culture with snGD2 or snCD19 J76 T cells. For some conditions, 10µg/ml neutralizing anti-INF−γ antibody was added to the co-cultures. **(B)** Surface expression of HLA-ABC on SKNDZ cells after 48h incubation with supernatants from snGD2 and snCD19 co-cultures show in panel A as determined by flow cytometry. **(C)** Changes in HLA-A2 mRNA expression in SKNDZ cells after 48h co-culture with synNotch J76 T cells as determined by quantitative RT-PCR. Data represent mean± SD from technical replicates (N=3). Statistical significance of differences between treatment conditions was determined by two-sided Student’s *t* test. **(D)** Expression of HLA-ABC on the surface of SKNDZ cells after 48h co-culture with snGD2 or snCD19 J76 T cells and subsequent removal of snGD2/snCD19 cells by cell sorting. Surface expression was determined after staining with an anti-HLA-ABC/APC antibody by flow cytometry. **(E)** NSG mice were subcutaneously injected with MYCN+ Kelly neuroblastoma cells and, after tumors reached a volume of 150mm^3^, received a single intratumoral injection of snGD2 or snCD19 J76 T cells. After 48h tumors were harvested and analyzed by HLA-ABC staining using immunohistochemistry. Data are representative of 2 biological replicates. **(F)** IFN-γ levels in the peripheral blood of NSG mice treated with snGD2 or snCD19 as determined by ELISA. Data represent mean ± SD from biological replicates (N=2). Statistical differences were determined by two-sided Student’s *t* test. **(G)** PRAME, NY-ESO-1 and housekeeping gene GAPDH expression on neuroblastoma cell lines as determined by RT-PCR. Data are representative of two independent experiments. **(H)** Cytotoxicity of NYESO1-or PRAME-specific TCR-transgenic or mock transduced primary human T cells against SKNDZ cells transduced with an NYESO1 expression construct, as determined by luminescence-based cytotoxicity assay. SKNDZ-ESO cells had been pretreated for 48h with primary snGD2i or snCD19i T cells. Data represent mean ± SD from technical replicates (N=3). Statistical significance was determined by two-sided Student’s *t*-test. **(I)** Schema of mouse model to determine the efficacy of primary snGD2 T cells together with PRAME-specific HSS1 T cells against neuroblastoma. **(J)** SKNDZ tumor volumes in mice treated with snGD2 or snCD19 CAR T cells together with HSS1 T cells. Data represent mean ± SD from individual animals per group (*N*=6). Statistical significance was determined by two-sided Student’s *t*-test. **(K)** SKNDZ cells were treated with snGD2i cells or snCD19i cells with or without addition of 10 ug/ml of an IFN-γ blocking antibody. SKNDZ cells were sorted by flow cytometry and mRNA expression analyzed by quantitative RT-PCR. Data represent mean ± SD from technical replicates (N=3). Statistical significance was determined by two-sided Student’s *t*-test. **(L)** SKNDZ cells were co-cultured with snGD2i cells or snCD19i cells for 48h and T cells were analyzed for expression of PD-1 by flow cytometry. Data represent a representative result from 2 independent experiments. **(M)** Schema of mechanism of action of snGD2i cells.

Finally, we explored additional effects of IFN-γ secreted by snGD2i cells on neuroblastoma and T cells. In addition to HLA itself, various proteins are required to generate peptide-loaded HLA, including β_2_-microglobulin (B2M), tapasin-1 (TAP1) and tapasin-2 (TAP2), as well as the various components comprising the immune proteasome, including LMP-2 and LMP-7 ^18,20,29^. It has previously been shown that IFN-γ can affect expression of many different proteins involved in immune recognition, and we therefore explored whether snGD2i treatment would also affect expression of any of the other proteins required for HLA peptide loading and presentation. We found that snGD2i treatment resulted in the significant upregulation of several components of the antigen processing machinery, including B2M, TAP1, and LMP2 (Fig. 4K) potentially in part contributing to T cell-mediated killing of neuroblastoma cells. In addition to its pro-inflammatory functions, IFN-γ has previously also been shown to induce the expression of immune checkpoints and significantly inhibit tumor-specific immune responses (*24*). Although we did observe increased T cell reactivity following snGD2i co-culture (Fig. 4H), indicating that treatment shifted the overall balance towards tumor cell killing, we next asked the question whether similar to HLA, immune checkpoints and/or its ligands would be upregulated following treatment. Surprisingly, we did not observe upregulation of immune checkpoints including PD-1 (Fig. 4L) and Tim-3 (Suppl. Fig. 4) on snGD2i cells themselves or PD-L1 on the neuroblastoma cells (Suppl. Fig. 5), indicating that while IFN-γ secreted by snGD2 cells was sufficient to restore HLA expression it was too low to induce immune checkpoints on neuroblastoma cells.

Taken together, we show here that snGD2i cells can be manufactured efficiently and specifically secrete IFN-γ when encountering GD2 positive neuroblastoma cells (Fig. 4M). The secreted IFN-γ in turn induces upregulation of HLA and other components of the antigen processing machinery but not immune checkpoints, facilitating substantial tumor cell killing by cancer antigen-specific T cells *in vitro* and *in vivo*.

## DISCUSSION

HLA loss represents one of the most common immune escape mechanisms in adult ^30^ and pediatric cancers ^1^. HLA loss not only renders the tumor cells invisible to the patient’s own immune system but also to adoptive TCR-based cell therapies. Non-genetic HLA loss is especially frequent in children with high-risk neuroblastoma ^31,32^, the most common solid extracranial tumor in childhood with an estimated 5-year overall survival of only 50% ^3^. We hypothesized that its reversal using neuroblastoma-sensing T cells delivering a customized IFN-γ payload may facilitate anti-tumor activity by restoring spontaneous T cell-mediated anti-neuroblastoma immunity, analogous to treatment with immune checkpoint inhibitors enabling pre-existing T cells to target tumor cells.

Previous clinical attempts to restore HLA expression to harness pre-existing tumor-specific T cell immunity using recombinant IFN-γ were largely unsuccessful ^33^. While few data are available regarding potential causes for those findings, it is likely that intratumoral concentrations of systemically injected IFN-γ, which has a short half-life of only 30 minutes ^34^, were too low to robustly induce HLA. Alternatively, it has been proposed that the immunosuppressive functions of IFN-γ, such as its ability to induce immune checkpoints on T cells and their ligands on tumor cells^35^, may have outweighed its pro-inflammatory effects.

We show here that IFN-γ delivered by snGD2i cells specifically induces HLA class I on neuroblastoma cells, sensitizing them to efficient killing by both PRAME and NYESO1-specific T cells after pretreatment with snGD2i cells. In contrast, we did not observe induction of immune checkpoints or their ligands. While we cannot rule out that the latter finding may at least in part be artificial and related to the assays used in our study, our data suggest that it may be possible to separate the deleterious effects of IFN-γ treatment from its benefits, possibly through timing, use of lower IFN-γ concentrations, or cellular context.

GD2 is not only expressed on neuroblastoma cells but also in the cerebellum and on peripheral nerves, resulting in pain-related side effects of GD2 CAR T cells currently in clinical trials and the two FDA-approved monoclonal GD2-specific antibodies Dinutuximab^36^ and Naxitamab^37^. However, as snGD2i cells do not exert any direct cytotoxic action, it is likely that side effects will be substantially milder than those of the other currently available GD2-targeted therapies. Similarly, side effects commonly observed during systemic pro-inflammatory treatment, such as checkpoint inhibition, will likely be less pronounced during snGD2i treatment due to the targeted delivery of IFN-γ to tumor cells and few healthy tissues. However, adverse events, such as tumor lysis and cytokine release syndrome, which commonly occur during treatment with all FDA-approved CAR T cell products and are the result of a strong immune-mediated anti-tumor response, may also occur during snGD2i treatment.

SynNotch technology used for the development of snGD2i cells has revolutionized the engineering of complex cell behavior in response to extracellular cues, such as chimeric antigen receptor logic gates and guided tissue differentiation ^23,24,38^. We here provide a proof-of-concept for another type of synNotch-based applications: the targeted delivery of pro-inflammatory cytokines to restore anti-tumor immunity. A key challenge for the clinical use of synNotch receptor-based cell therapies remains the potential immunogenicity resulting from the use of non-human transcription factors needed to ensure orthogonal gene regulation, which may result in limited persistence. While efforts to develop human transcription factor-based synNotch systems are underway to address this concern ^39,40^, our finding that HLA induction is relatively stable even after removing snGD2i cells suggests that long-term persistence may not always be necessary. A key consideration for synNotch T cells that lack a conventional T cell activation component is the potential impact on their local expansion, possibly resulting in relatively low intratumoral snGD2i numbers. In this study we performed intratumoral T cell injections and therefore were not able to investigate this potential drawback of this approach. If reduced local snGD2i expansion does in fact limit their efficacy *in vivo*, a potential solution to this could be the combination of synNotch signaling components with a transgenic tumor-specific TCR in the same cell or to combine the IFN-γ construct with an IL-2 and IL-2Rγ expression cassette to simulate T cell activation upon target recognition ^41^.

Taken together, we provide a preclinical proof-of-concept for the targeted induction of HLA on cancer cells restoring their sensitivity to T cell-mediated tumor cell killing. This opens the door to the development of novel, highly specific and likely safe treatment approaches for many common types of cancer showing reversible HLA loss.

## AUTHOR CONTRIBUTIONS

F.I. and T.L. conceived the project, planned, and performed experiments, analyzed data, and wrote the manuscript. S.W., H.Z., S.Y., S.V.A., E.M., and M.L.O. performed experiments and analyzed data. M.F., S.V.R., and S.M. analyzed data.

## ACKNOWLEDGEMENTS

This project was supported by a St. Baldrick’s Foundation Fellowship (F.I.), an NCCN Young Investigator Award (T.L.), a University of Utah Center for Clinical & Translational Science (CCTS) award (T.L.), and the Huntsman Cancer Institute Experimental Therapeutics program supported by the National Cancer Institute of the National Institutes of Health under Award number P30CA042014. (D.A. and T.L.). We thank the Huntsman Cancer Institute in Salt Lake City, UT for the use of the Preclinical Research Resource (PRR), which performed the neuroblastoma xenograft model, and the University of Utah and University of Maryland Flow Cytometry core facilities, which assisted with flow cytometry analyses and cell sorting. pHIV-Luc-ZsGreen was a gift from Bryan Welm (Addgene #39196) and SFG.CNb30_opt.IRES.eGFP was a gift from Martin Pule (Addgene # 22493). pHR_Gal4UAS_IRES_mC_pGK_tBFP (Addgene # 79123), pHR_Gal4UAS_tBFP_PGK_mCherry (Addgene # 79130), and pHR_PGK_antiCD19_synNotch_Gal4VP64 (Addgene # 79125) were gifts from Wendell Lim. The PRAME-specific HSS1 TCR was a gift from Mirjam HM Heemskerk (University Medical Center Leiden, Netherlands).

## METHODS

### Cell lines and primary human cells

Cell lines SKNDZ, LAN5, SHSY5Y, SKNFI, SKNSH, NB1643, Kelly, and Phoenix-Ampho cells were purchased from ATCC and cultured according to ATCC instructions. Lenti-X 293T cells were purchased from Takara and cultured according to the manufacturer’s instructions. J76 cells were a kind gift by Dr. Uckert (Max Delbrueck Center for Molecular Medicine, Germany) and cultured in RPMI-1640 (ATCC formulation) supplemented with 10% fetal bovine serum. To determine the killing of neuroblastoma cells by transgenic T cells, SKNDZ cells were transduced with pHIV-Luc-ZsGreen lentivirus as well as a lentiviral NYESO1 expression construct carrying a BFP reporter and sorted on a FACSaria flow cytometer (BD) for GFP and BFP expression. Healthy donor buffy coats were obtained from the Blood Centers of America and the New York Blood Center and peripheral blood mononuclear cells were isolated from buffy coats by density gradient using FicollPaque (GE) as previously described (*25*). Tissue sections of de-identified primary neuroblastoma tumor samples were obtained from Dr. Joshua Schiffmann at the University of Utah.

### Quantitative RT-PCR

Expression of HLA class I, HLA-A2, NYESO1, PRAME, as well as antigen processing components was measured by quantitative reverse transcription PCR. When analyzing neuroblastoma cells from co-cultures with primary snGD2i cells, neuroblastoma cells were first purified by cell sorting using a FACSaria flow cytometer (BD). Total RNA was isolated using the RNeasy Mini kit (Qiagen), according to the manufacturer’s instructions. Complementary DNA (cDNA) was generated using the SuperScript III First-Strand Synthesis system (Thermo-Fisher). Quantitative PCRs were performed on a CFX96 Real Time PCR system (Bio-Rad) using the primers listed in Supplementary Table 2 using SsoFast EvaGreen Supermix (Bio-Rad). The expression of the respective gene of interest was normalized to the samples’ respective GAPDH expression using the ΔCt or ΔΔCt method as indicated in the figures. For co-cultures with snGD2i cells based on the HLA-A2-negative cell line J76, neuroblastoma cells were not sorted but HLA-A2 expression normalized to CD171 expression.

### SynNotch and TCR T cell production

Previously described lentiviral transfer constructs pHR-PGK-antiCD19-synNotch-Gal4VP64 and pHR-Gal4UAS-IRES-mC-PGK-tBFP were obtained from Addgene. The PRAME-specific HSS1 TCR construct was provided by Dr. Heemskerk (Leiden University Medical Center, Netherlands) and confirmed by Sanger sequencing. The NYESO1-specific 1G4-LY TCR construct was synthesized by Twist Bioscience and cloned into the pRRL-cPPT-PGK-GFP-WPRE backbone as previously described. GD2-specific scFv sequences for clones KM666, 3F8 VLVH, 3F8 VHVL, 5F11, and 14.18 were synthesized by Twist Bioscience and used to replace the CD19-specific scFv using splice-by-overlap-extension. Similarly, the open-reading frame of full-length IFN-γ was synthesized by Twist Bioscience and cloned into pHR-Gal4UAS-IRES-mC-PGK-tBFP in front the IRES. To generate primary T cells, the synNotch receptor construct was also transferred into the gammaretroviral SFG backbone. Lentivirus was generated by co-transfection of Lenti-X 293T cells with lentiviral transfer plasmids, pMD2.G (VSV-G envelope), and psPAX2 using Lipofectamine 2000 according to the manufacturer’s instructions. Amphotropic gammaretrovirus was generated by transfection of Phoenix-Ampho cells (ATCC # CRL-3213) using Lipofectamine 2000 according to the manufacturer’s instructions. Virus-containing supernatants were concentrated with Lenti-X and Retro-X concentrator (Takara), respectively, and transferred to Retronectin-coated 24 well plates on the day of transduction. PBMCs were stimulated for 2 days with CD3/CD28 T cell activation beads (Thermo # 11131D) in the presence of 40IU/mL IL2 (R&D Systems # 202-IL-010) in AIM V (Thermo) supplemented with 5% human serum (Sigma #H3667) and incubated at 37°C/5% CO_2_. Bead-stimulated cells were transferred to Retronectin-coated (Takara) virus-containing plates and incubated overnight. Transduction was repeated the next day before counting and diluting cells to 0.4×10^6^ cells/ml. After the second transduction cells were grown for an additional 7 days before removing beads using a DynaMag-15 magnet (Thermo). IL-2 was replenished every 2 days to 40IU/mL. Cells were frozen in 90% FCS/10% DMSO and stored in liquid nitrogen.

### Flow cytometry analysis

Flow cytometry staining and analyses were performed as previously described (*25*). Antibodies used for flow cytometry analyses are listed in Supplementary Table 1. Commercially available antibodies were used at dilutions recommended by the respective manufacturer. Flow cytometry data was acquired on an LSR Fortessa or LSR II flow cytometer (BD) and analyzed using FlowJo 10 (BD).

### HLA induction and cytotoxicity co-cultures

To determine induction of HLA and immune checkpoints, as well as sensitization of neuroblastoma cell lines by snGD2 cells, 2.5×10^5^ luciferase expressing neuroblastoma cells were seeded in each well of a flat bottom 6 well plate. After 3-4h, the indicated ratio of synNotch T cells was added, and the co-culture was incubated for 72h at 37C/5% CO2. To determine killing of neuroblastoma cells by TCR transgenic T cells following synNotch T cell co-culture, co-cultures were washed twice with PBS to remove synNotch T cells and neuroblastoma cells were subsequently detached using trypsin. 5×10^4^ target cells/well were plated in a black 96 well plate and the indicated ratio of TCR-transgenic T cells was added to the tumor cells. Cells were co-cultured overnight and luminescence was measured the next morning after addition of 150 μg/ml D-luciferin (Gold Biotechnology Cat# LUCNA-2G) and subsequent incubation for 5 mins at 37°C. Luminescence was determined on a multi-mode plate reader (Tecan Spark).

### Cytokine detection

Supernatants were harvested from 96 well overnight co-cultures and immediately frozen at -80°C. IFN© concentrations were determined via standard curve using a commercial ELISA kit according to the manufacturer’s instructions (Thermo). Absorbance was measured on a multi-mode plate reader (Tecan).

### Mouse xenograft study

7-8-week-old female NSG mice were randomly grouped and sublethally irradiated at 5cGy on the morning of day -2 followed by subcutaneous administration of 2.5×10^5^ SKNDZ-ESO-Luc. Once tumors reached a size of 100mm^3^, 5×10^6^ snGD2i or snCD19i cells were injected intratumorally. Mice were sacrificed 72h after T cell injection and tumors embedded in paraffin prior to IHC analysis. To determine the anti-tumor activity of snGD2i cells together with TCR-transgenic T cells, 7-8-week-old female NSG mice were randomly grouped and sublethally irradiated at 5cGy on the morning of day -2 followed by subcutaneous administration of 3×10^6^ SKNDZ-ESO-Luc together with 5×10^6^ snGD2i or snCD19i cells and 5×10^6^ HSS1 TCR T cells. Tumor size was determined by caliper measurements. Animal work was performed by the Preclinical Research Resource at the University of Utah and the Translational Shared Service at the University of Maryland.

### Immunohistochemistry staining of neuroblastoma tumors

Immunohistochemistry staining of paraffin-embedded tumor sections from patients with neuroblastoma was performed using standard procedures. First, sections were deparaffinized using a series of Citrisolv (Fisher # 22-143-975)/ethanol dilutions and unmasking was performed using a citrate-based unmasking solution (Vector # H-3300-250). Slides were washed and endogenous peroxidase was blocked using 3% hydrogen peroxide. Subsequently, slides were blocked in PBS containing 10% BSA, 10% human serum (Sigma-Aldrich), 10% goat serum (Jackson Immuno), 0.5% Tween-20, avidin/biotin blocking solution (Vector Laboratories # SP-2001), mouse (Miltenyi Biotec # 130-092-575) and human (Miltenyi Biotec # 130-059-901) FcR blocking reagents, and M.O.M. blocking solution. Primary HLA class I-specific antibody (clone W6/32, Biolegend # 311428) was added at a dilution of 1:50 and slides were incubated in a humidity chamber over night at 4C. The next day, slides were developed using the M.O.M. Immunodetection kit (Vector Laboratories # BMK-2202), ABC HRP kit (Vector Laboratories # PK-4000), and the DAB peroxidase substrate kit (Vector Laboratories # SK-4100) according to the manufacturer’s instructions.

### Statistical considerations

Significance of differences in mean fluorescence intensity, mRNA levels, cytokine concentrations, and tumor cell viability were calculated by Student’s t-test. All statistical tests were performed using Prism 7 (GraphPad Software). Results were considered significant when *p* <0.05. * *p* <0.05, ** *p* < 0.01, *** *p* < 0.001, *p* < 0.0001.

### Study approval

Informed consent was obtained from all patients and samples were collected with the approval of the Institutional Review Board of the University of Utah (IRB # 49682, Principal Investigator: Dr. Joshua Schiffmann). Animal procedures were conducted under protocol #16-05007, approved by the Institutional Animal Care and Use Committee at the University of Utah or under protocol #1021001, approved by the Institutional Animal Care and Use Committee at the University of Maryland Baltimore.

