## Supplementary Material for "Targeted delivery of interferon-gamma by synNotch T cells sensitizes neuroblastoma cells to T cell-mediated killing"

### SUPPLEMENTARY FIGURES

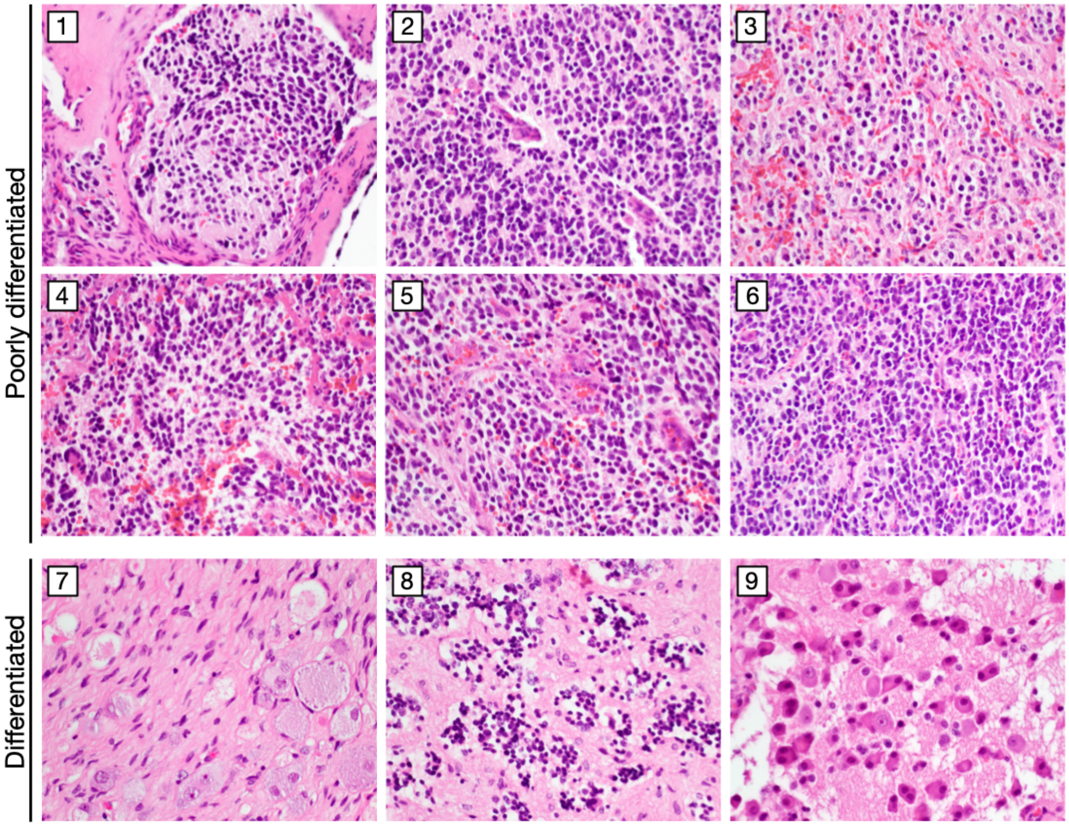


**Supplementary Figure 1: H&E Staining of primary neuroblastoma tumor samples.**

Microscopy images of primary neuroblastoma samples shown in Fig. 1 after staining with hematoxylin and eosin (H&E). 1-6: Poorly differentiated neuroblastomas (low-intermediate MKI), original magnification 100x. 7-8: Ganglioneuroblastoma, intermixed subtype, original magnification 200x. 9: Differentiating neuroblastoma.

| Cell line | HLA-A | RPKM |
|---|---|---|
| Kelly | 01:01<br>01:01 | 28.37 |
| SKNDZ | 24:03<br>02:01 | 12.86 |
| SKNFI | 23:01<br>03:01 | 23.52 |
| SKNSH | 24:02<br>01:01 | 60.99 |

**Supplementary Figure 2: HLA-A alleles and mRNA expression in neuroblastoma cell lines.** Each cell line's respective HLA-A alleles and HLA-A mRNA expression in reads per kilobase per million reads (RPKM) were obtained from the TRON Cell Line Portal [1].

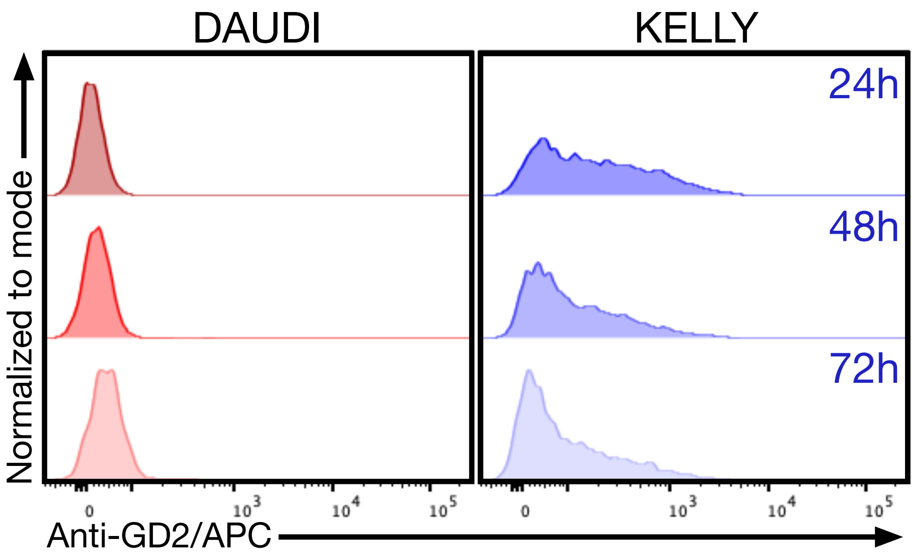


**Supplementary Figure 3: Detection of GD2 on the surface of snGD2i cells after co-culture.** GD2 was analyzed on primary human snGD2i cells after indicated durations of co-culture with GD2-negative lymphoma cell line Daudi or the GD2+ neuroblastoma cell line Kelly. Co-cultures were stained using a monoclonal GD2-specific antibody and analyzed by flow cytometry. Plots show results from at least 3 independent experiments.

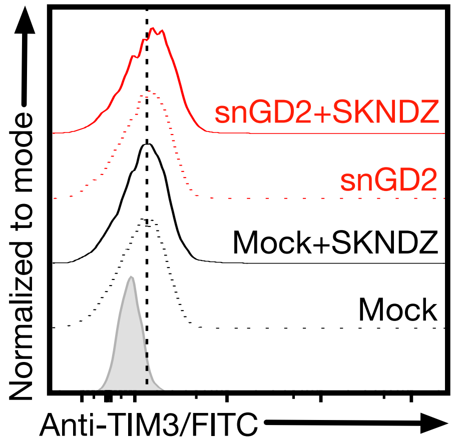


**Supplementary Figure 4: Lack of induction of TIM-3 on snGD2i cells following co-culture with SKNDZ cells.** SKNDZ cells were co-cultured with snGD2i cells or snCD19i cells for 48h and T cells were analyzed for expression of TIM-3 by flow cytometry. Data are a representative result from 2 independent experiments.

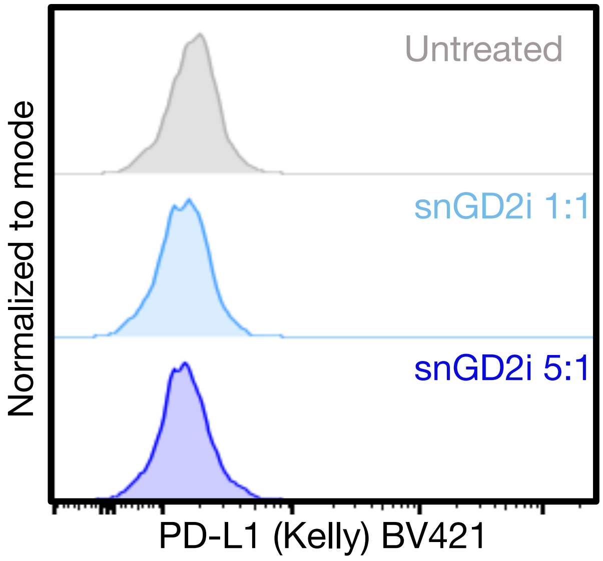


**Supplementary Figure 5: Lack of induction of PD-L1 on neuroblastoma cells following co-culture with snGD2i cells.** Kelly neuroblastoma cells were co-cultured with snGD2i cells at different effector-target ratios for 72h and Kelly cells were analyzed for expression of PD-L1 by flow cytometry. Data are a representative result from 2 independent experiments.
